# Depiction of secondary metabolites and antifungal activity of *Bacillus velezensis* DTU001

**DOI:** 10.1101/577817

**Authors:** Sagarika Devi, Heiko T. Kiesewalter, Renátó Kovács, Jens Christian Frisvad, Tilmann Weber, Thomas Ostenfeld Larsen, Ákos T. Kovács, Ling Ding

## Abstract

For a safe and sustainable environment, effective microbes as biocontrol agents are in high demand. We have isolated a new *Bacillus velezensis* strain DTU001, investigated its antifungal spectrum, sequenced its genome, and uncovered the production of lipopeptides in HPLC-HRMS analysis. To test the antifungal efficacy, extracts of *B. velezensis* DTU001 was tested against a range of twenty human or plant pathogenic fungi. We demonstrate that inhibitory potential of *B. velezensis* DTU001 against selected fungi is superior in comparison to single lipopeptide, either iturin or fengycin. The isolate showed analogous biofilm formation to other closely related *Bacilli*. To further support the biocontrol properties of the isolate, coculture with *Candida albicans* demonstrated that *B. velezensis* DTU001 exhibited excellent antiproliferation effect against *C. albicans*. In summary, the described isolate is a potential antifungal agent with a broad antifungal spectrum that might assist our aims to avoid hazardous pathogenic fungi and provide alternative to toxicity caused by chemicals.

## 1. Introduction

The prerequisite for benign and effective antifungal agents increases in parallel with the existing and escalating emergence of fungal pathogens resistant to current therapies [1], being relevant to both therapeutics and biocontrol. While infectious diseases impose serious public health burden and often have devastating consequences, plant diseases constitute an emerging threat to global food security. Additionally, extensive use of chemicals to control plant diseases has distressed the ecological balance of microbes inhabiting soil. This in turn has led to the appearance of resistant pathogenic strains, groundwater contamination, and obvious health risks to human beings. Thus, the identification and development of environmental friendly alternatives to the currently used chemical pesticides for combating a variety of crop diseases is nowadays one of the biggest ecological challenges [2]. It is therefore vital to obtain a broad range of antifungal agents that are non-hazardous to human health as well as to the environment.

Several microorganisms have so far been discovered for the development of biopesticides at commercial level. Among the various microorganisms, *Bacillus* spp., *Pseudomonas* spp., and some *mycorrhizal fungi* are widely used groups for biocontrol [3]. Especially, the *Bacillus* genus was proven to be source an excellent source of antifungal agents and thus *Bacilli* are widely used as biocontrol agents [4,5]. The US Food and Drug Administration (US FDA) declared *Bacillus subtilis* as GRAS (generally recognized as safe) organisms for its use in food processing industries [6]. Members of the *B. subtilis* groups were found to produce numerous highly active antifungal compounds [7]. The *B. subtilis* group contains the “Operational Group *Bacillus amyloliquefaciens*” that includes *B. amyloliquefaciens*, *Bacillus velezensis*, and *Bacillus siamensis* [8]. Among these, *B. velezensis* FZB42 (formerly *B. amyloliquefaciens* subsp. *plantarum* FZB42) is the most researched commercially used *Bacillus*-based biofertilizer and biocontrol agent in agriculture [9].

Antifungal cyclopeptides appeared to play a major role in biological control of plant pathogens [4, 10]. Composed of a peptide and a fatty acyl moiety, these amphiphilic metabolites have consolidated their position as potent therapeutic compounds, such as daptomycin, a lipopeptide antibiotic used in human medicine [11]. Antibacterial, -viral, -fungal, -tumor activity and immune-modulatory effects are some of their proven and emerging properties [12]. In addition, secondary metabolites seem to play an important role in bacterial-fungal interaction of *Bacilli* [13] and other bacterial life styles, including surface spreading [14, 15, 16].

Here, we describe a new *Bacillus* isolate DTU001, which exhibited a broad and strong antifungal spectrum against different pathogenic fungi ranging from plant pathogens to human pathogens. Using *in silico* DNA hybridization, it was identified as *B. velenzensis*. To dissect the genetic potential as an antifungal agent, we genome-mined BGCs in 31 *Bacillus* species and compared their biosynthetic potential with a special focus on commercially applied strains. To detect the synthesized secondary metabolites of *B. velezensis* DTU001, the lipopetide profile of the strain’s crude extract was investigated.

Lipopeptides produced by *Bacilli*, such as iturin, fengycin and surfactin are believed to contribute to their antifungal potential. When new isolates from the *B. subtilis* species complex are evaluated, the role of single lipopeptide in antifungal activity is scrutinized lacking systematic *in vitro* evaluation of the antifungal efficacy of the combination of these secondary metabolites. To fill in this gap, we also assayed the efficacy of extracted iturin and fengycin in comparison to the crude extract of *B. velezensis* DTU001 using twenty critical human and plant pathogens. In this report, we outline the phylogeny, genomics, metabolomics, biofilm formation, and antifungal effects of *B. velezensis* DTU001.

## 2. Materials and methods

### 2.1. Bacterial and fungal strains

*B. velezensis* DTU001 is an airborne bacterium, isolated indoor at the DTU campus, Kongens Lyngby Denmark. Twenty selected plant/human pathogens (Table 1) covering five genera *Aspergillus*, *Penicillium*, *Talaromyces*, *Cryptococcus* and *Candida* were obtained from the DTU Bioengineering IBT fungal collection. All microbes are maintained at the department.

**Table 1.**
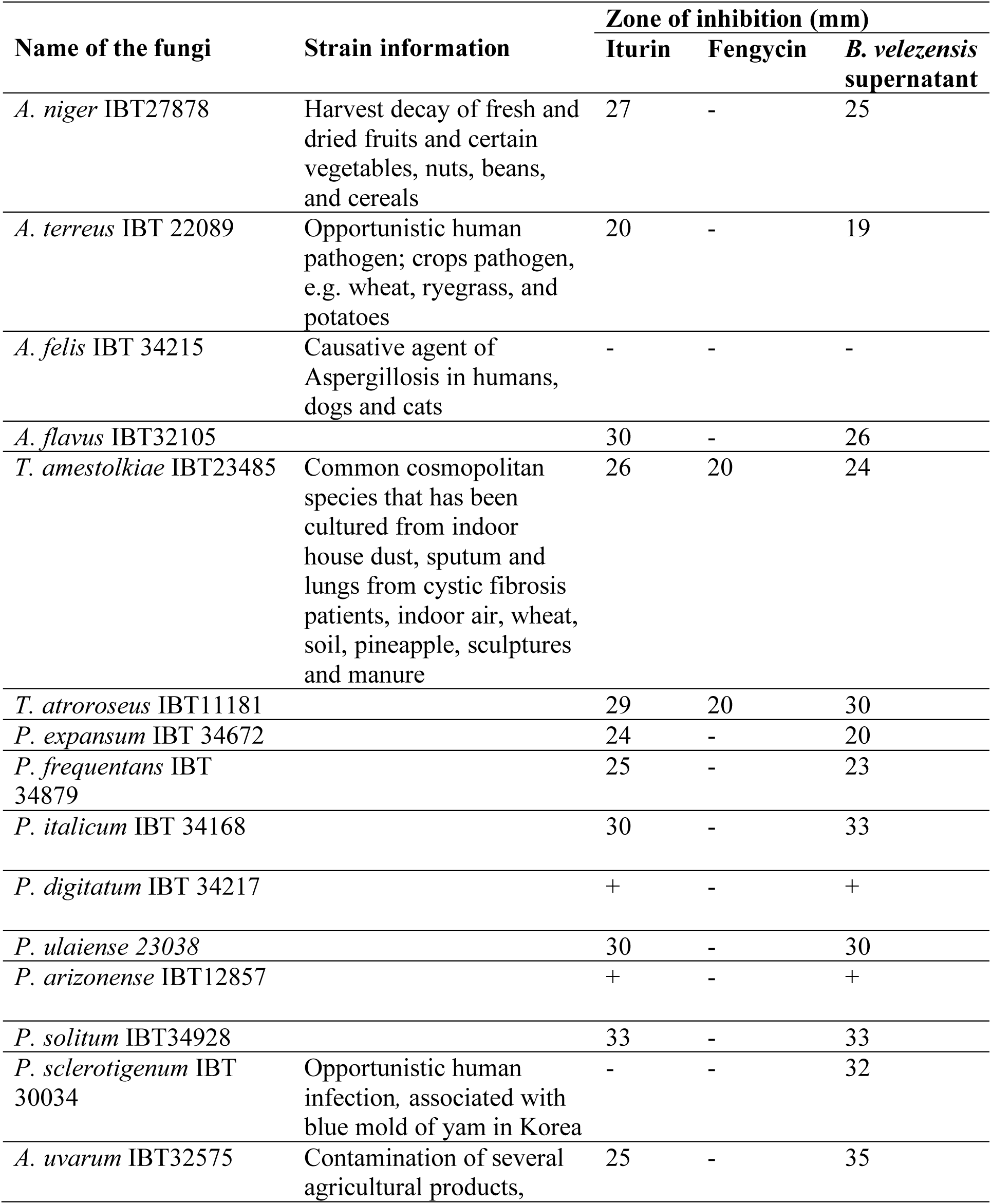
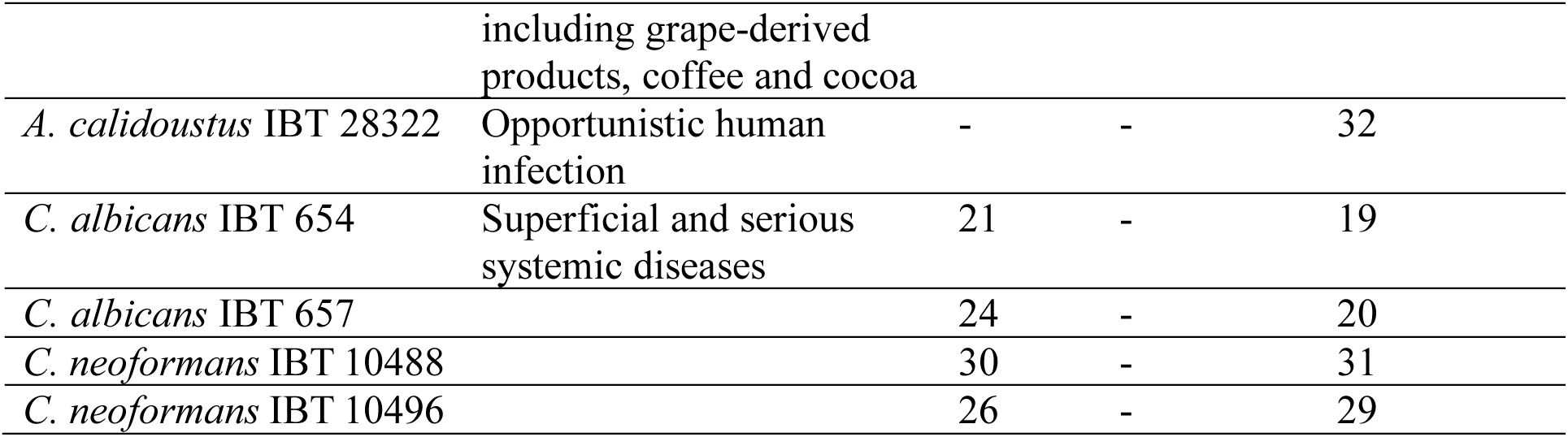
Bioactivity of iturin (1mg/mL), fengycin (1mg/ mL), and crude *B. velezensis* extract (1mg/ mL) against 16 human- and plant-pathogenic filamentous fungi and 4 yeasts

### 2.2. Genome sequencing

Genomic DNA was isolated from *B. velezensis* DTU001 using Bacterial and Yeast Genomic DNA kit (EURx, Poland). Genome sequencing has been performed on PacBio RSII platform, *de novo* assembly and annotation were done using the RS HGAO Assembly (v3.0) and Prokka (v1.12b) software, respectively. The complete sequence has been deposited in NCBI (accession number CP035533) followed by automatic annotation via NCBI PGAP [17].

### 2.3. Chemicals

Solvents used in this study were LC–MS grade, and all other chemicals were analytical grade. Iturin A, fengycin and solvents as well were from Sigma-Aldrich (Steinheim, Germany) unless otherwise stated. Water was purified using a Milli-Q system (Millipore, Bedford, MA). ESI– TOF tune mix was purchased from Agilent Technologies (Torrance, CA, USA).

### 2.4. Digital DNA–DNA hybridization (dDDH)

The genome-to-genome-distance calculator (GGDC) version 2.1 provided by DSMZ (http://ggdc.dsmz.de/) was used for genome-based species delineation [18] and genome-based subspecies delineation [19]. Distances were calculated by (i) comparing two genomes using the chosen program to obtain HSPs/MUMs and (ii) inferring distances from the set of HSPs/MUMs using three distinct formulas. Next, the distances were transformed to values analogous to DDH. The DDH estimates were based on an empirical reference dataset comprising real DDH values and genome sequences. The DDH estimate resulted from a generalized linear model (GLM) which also provided the estimate’s confidence interval (after the ± sign). Three formulas are available for the calculation: (1) HSP length/total length; (2) identities/HSP length; and (3) identities/total length. Formula 2, which is especially appropriate to analyze draft genomes, was used in this study.

### 2.5. Construction of the evolutionary tree

An initial whole genome phylogeny was inferred using the automlst tool (http:/automlst.ziemertlab.com; Ziemert et al., personal communication). This automatically identifies related strains based on a seed collection of sequences obtained from NCBI Genbank (*B. velezensis* DTU001, *B. thuringiensis* serovar Berliner ATCC 10792, *B. cereus* 03BB102, *B. megaterium* DSM319, *B. velezensis* FZB42), using a MASH-based algorithm, that extracts 80 phylogenetically relevant genes and builds a MLST phylogeny out of these data. The sequence alignment obtained from automlst was then manually refined (removal of very similar species). To infer the best phylogenetic model, jModeltest (v.2.1.10) was used with default parameters [20]. In all three evaluation criteria (Akaike Information Criterion AIC, Bayesian Information Criterion BIC and the Decision Therory Performance-based Selection), the GTR+I+G model was deemed most suitable. Thus, a maximum-likelihood tree was constructed in PhyML (v3.3.20180621) [21] using the parameters suggested by jModeltest. The tree then was checked with Dendroscope (v3.5.10) [22] and plotted using the ETE3 toolkit [23].

### 2.6. Genome mining of secondary metabolites

*B. velezensis* DTU001 and other 32 Bacillus strains were mined for the biosynthetic gene clusters using the antiSMASH platform [24] and BiG-SCAPE [25].

### 2.7. Agar-diffusion assay

Agar-diffusion assay was carried to test the anti-fungal activities. *B. velezensis* DTU001 grown on agar plates was extracted by ethyl acetate, concentrated and re-dissolved in methanol (1 mg/mL). DMSO was used as a negative control. Initial screening of the bacterial extract was done by standard agar disc diffusion method [26] and the zone of inhibition was noted. The 16 pathogenic filamentous fungi were grown on sterile MEAOX plates and the 4 yeasts were cultivated on PDA at 25 ^º^C. 5-days old fungal plates and 2-days old yeast plates were taken for further investigation. Spore suspensions harvested in tween 20 from MEAOX medium plates with a final cell concentration of 5×10^5^ cfu/mL of each microbial culture (fungi and yeast) were maintained for the disc diffusion assay. Plugs were harvested from 48 hr grown *B. velezensis* DTU001 from MEAOX plates and extracted in isopropanol-ethyl acetate-1% formic acid followed by 1 hr ultra-sonication. The supernatant was collected and dried utilizing nitrogen evaporator. The test sample (*B. velezensis* extract) and the standard compounds (iturin and fengycin) were dissolved in 0.5% DMSO to a concentration of 1 mg/mL each. Sterile Muller Hinton Agar plates were inoculated with the test organisms using lawn culture method. Discs of 6mm were cut at the centre of the sterile plates using a sterile cork borer. The discs were loaded with 50 µl of each test sample. The plates were maintained at 25 ± 2 ^º^C for 5 days and observed for clear zone of inhibition. Iturin and fengycin standards were maintained as positive control. Blank assays in DMSO were applied to avoid any possible effect of the solvent. Each experiment was conducted in triplicates for further confirmation and reproducibility of results.

### 2.8. Anti-Candida effects under planktonic and biofilm conditions

The impact of *B. velezensis* DTU001 on *C. albicans* SC5314 reference strain was assayed. *C. albicans* and *B. velezensis* were subcultured on Sabouraud dextrose agar and lysogeny broth (LB) agar, respectively. The growth of the individual strains in both monoculture and coculture was evaluated in order to examine the effect of *B. subtilis* on *C. albicans* planktonic growth. The final cell concentrations were adjusted to 2×10^5^ cells/ml in RPMI-1640 broth + LB broth (50:50 v/v %) both for *C. albicans* and *B. subtilis*. Test tubes were incubated with agitation in darkness at 35 °C. 100 µl culture was removed at 0, 2, 4, 6, 8, 12 and 24 hours, serially diluted tenfold in sterile physiological saline then plated onto either Sabouraud dextrose agar or LB agar. Plates were incubated for 24 and 48 hours at 35 °C for *B. subtilis* and *C. albicans*, respectively. Results are representative of three independent experiments and are expressed as mean ± SD (error bars), which was presented using GraphPad Prism 6.05.

For the biofilm assays, polystyrene flat-bottom 96-well microtiter plates (TPP, Trasadingen, Switzerland) were used for the formation of single and mixed species biofilms of *C. albicans* and *B. subtilis*. A volume of 200 µl of cell suspension (1×10^6^ cells/mL in RPMI-1640 broth + LB broth (50:50 v/v%) for *C. albicans* and 1×10^6^ cells/mL in RPMI-1640 broth + LB broth (50:50 v/v%) for *B. subtilis*) was pipetted to each well for single biofilms, while 100 µL from each suspension (2×10^6^ cells/mL for *C. albicans* with 2×10^6^ cells/ml for *B. subtilis*) was added for mixed species biofilms to wells assigned to endpoints 2, 4, 6, 8, 12 and 24 hours. The plates were incubated in darkness with static conditions. After 2, 4, 6, 8, 12 and 24 hours of incubation, the corresponding wells were washed three times with sterile physiological saline then 200 µL sterile saline was added into wells, scraped off the adhered cells from surfaces; afterwards, aliquots of 100 µL were removed. Cells were released by mild sonication and vortexing then diluted serially ten-fold and plated onto either Sabouraud dextrose agar or LB agar. Plates were incubated for 24 and 48 hours at 35 °C for *B. subtilis* and *C. albicans*, respectively. Results are representative of three independent experiments and are expressed as mean ± SD (error bars), which was presented using GraphPad Prism 6.05.

### 2.9. Bacterial biofilm formation

Bacterial strains (*B. velezensis* DTU001, *B. subtilis* NCBI 3610 [28] (Branda et al., 2001), *B. subtilis* ATCC6633 (ATCC collection), *B. subtilis* PS216 [27], *B. velezensis* FZB42 (obtained from Bacillus Genetic Stock Centre), and *B. methylotrophicus* NCCB100236 (obtained from Bacillus Genetic Stock Centre) were inoculated in lysogeny broth (Lenox LB, Carl Roth, Germany) and grown overnight for 16 hours at 37°C at 225 rpm. For pellicle development 20 μl was inoculated in 2 mL MSgg medium and placed in a 24-well plate, while 5 μL spotted on MSgg medium solified with 1,5 % agar that was prepared as described previously [8, 28]. Both pellicle and colony biofilms were incubated at incubated at 30°C static conditions and imaged after 3 days.

### 2.10. Secondary metabolites identification

A UHPLC–DAD–QTOF method was set up for screening, with typical injection volumes of 0.1–2 µL extract. Separation was performed on a Dionex Ultimate 3000 UHPLC system (Thermo Scientific, Dionex, Sunnyvale, California, USA) equipped with a 100×2.1 mm, 2.6 µm, Kinetex C_18_ column, held at a temperature of 40 °C, and using a linear gradient system composed of A: 20 mmol L^−1^ formic acid in water, and B: 20 mmol L^−1^ formic acid in acetonitrile. The flow was 0.4 mL min^−1^, 90 % A graduating to 100 % B in 10 min, 100 % B 10–13 min, and 90 % A 13.1–15 min.

Time-of-flight detection was performed using a maXis 3G QTOF orthogonal mass spectrometer (Bruker Daltonics, Bremen, Germany) operated at a resolving power of ~50000 full width at half maximum (FWHM). The instrument was equipped with an orthogonal electrospray ionization source, and mass spectra were recorded in the range *m*/*z* 100–1500 as centroid spectra, with five scans per second. For calibration, 1 µL 10 mmol L^−1^ sodium formate was injected at the beginning of each chromatographic run, using the divert valve (0.3–0.4 min). Data files were calibrated post-run on the average spectrum from this time segment, using the Bruker HPC (high-precision calibration) algorithm.

For ESI^+^ the capillary voltage was maintained at 4200 V, the gas flow to the nebulizer was set to 2.4 bar, the drying temperature was 220 °C, and the drying gas flow was 12.0 L min^−1^. Transfer optics (ion-funnel energies, quadrupole energy) were tuned on HT-2 toxin to minimize fragmentation. For ESI^−^ the settings were the same, except that the capillary voltage was maintained at −2500 V. Unless otherwise stated, ion-cooler settings were: transfer time 50 µs, radio frequency (RF) 55 V peak-to-peak (Vpp), and pre-pulse storage time 5 µs.

## 3. Results and discussion

### 3.1. The genome of the indoor airborne isolate B. velezensis DTU001 encodes numerous secondary metabolites

*B. velezensis* DTU001 was isolated indoor at the DTU campus in Denmark and selected for detailed analysis based on its potential antimicrobial activity. The complete genome sequence of *B. velezensis* DTU001 consisted of a circular 3.927.210 bp chromosome with a GC content of 46.6%. Its 16S rRNA gene sequence exhibited 99% similarity to *B. velezensis* and *B. amyloliquefaciens* in genome database.

The phylogenetic affiliation of DTU001 isolate was further investigated using digital DNA:DNA-hybridization (dDDH) similarities and differences in genomic G+C content. (https://ggdc.dsmz.de/). These distances are transformed to values analogous to DDH using a generalized linear model (GLM) inferred from an empirical reference dataset comprising real DDH values and genome sequences. Compared to *B. velezensis* FZB42 (NC-009725.1), DDH value of 80.3%, difference in % G+C: 0.10. Meanwhile, there is a DDH analogous value of *B. amyloliquefaciens* (FN597644.1) 55.4%, difference in % G+C: 0.51. This result clearly supports that DTU001 is closer related to *B. velezensis*, where the DDH value exceeded the threshold of 70%.

A phylogenetic analysis with automlst (http://automlst.ziemertlab.com) was carried out, which automatically extracts 80 conserved genes, concatenates them, computes a multiple sequence alignment and builds a phylogenetic tree. The sequence alignment was manually checked, and a maximum-likelihood tree refined with jModelTest2 [30] and PhyML [21]. According to the MLST analysis, *B. velezensis* DTU1 forms a clade with other *Bacilli* that were recently proposed to be assigned the new species *Bacillus lentimorphus* [29].

To determine the genetic potential of *B. velezensis* DTU001 as biocontrol strain, the complete genome was surveyed for biosynthetic gene clusters that identified nine gene clusters involved in nonribosomal synthesis of lipopeptides, polyketides, and siderophore bacillibactin. About 15.4 % of the whole DTU001 genome was devoted to secondary metabolites synthesis of nonribosomal synthesis of antimicrobial compounds and siderophores. In addition, genes for synthesis of the highly bioactive extracellular alkaline guanyl-preferring ribonuclease, previously detected in *B. amyloliquefaciens* [30] was also identified in the genome of DTU001. In order to compare the biosynthetic potentials between different *Bacilli*, *B. velezensis* together with other 17 species were mined for the genetic potential of secondary metabolite production using antiSMASH and BiG-SCAPE. All *Bacilli* related to *B. velezensis* DTU001 have the genetic potential to utilize bacillibactin as a siderophore to acquire irons from the environments (Fig. 1). All selected *B. amyloliquefaciens*, *B. velezensis*, and *B. subtilis* strains showed high similarities in BGC potentials and are considered as talented strains producing various lipopetides or polyketides. More distantly related *Bacillus* strains, such as members of the *Bacillus cereus* group showed a different secondary metabolite production genotype. All of these strains (*B. cereus*, *B. anthracis, B. thuringiensis*) seem to be unable of producing lipopeptides of the surfactin, iturin or fengycin families.

**Fig. 1.**
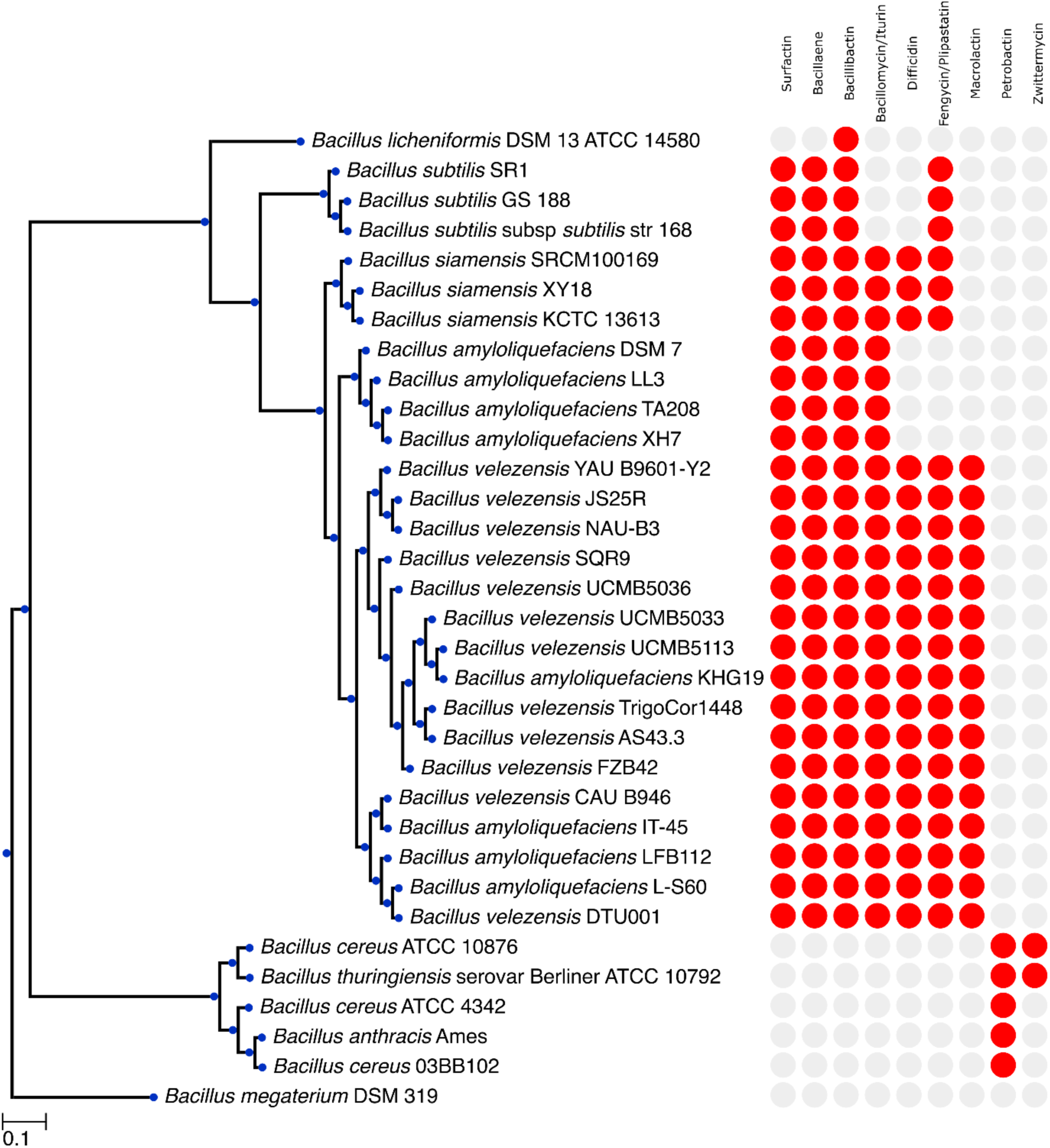
Evolutionary relationships of taxa between *B. velezensis* DTU001 and other *Bacilli*. A maximum-likelihood phylogenetic tree based on 80 conserved genes of selected strains of *Bacilli* is shown; *B. megaterium* DSM 319 was selected as outgroup. The presence/absence of typical *Bacillus* BGCs is depicted on the right.

### 3.2. Antifungal characterization of B. velezensis

Iturins are well-known secondary metabolites important for biocontrol against plant pathogens such as *Xanthomonas campestris* subsp. *Cucurbitae, Pectobacterium carotovorum* subsp*. Carotovorum, Rhizoctonia solani, Fusarium graminearum* etc. [31, 32]. However, the impact of iturins against medically-relevant fungal pathogens is less explored. Therefore, we tested the efficacy and target specificity of DTU001 against selected important plant and human pathogenic fungi. In addition, we compared the effectiveness of the DTU001 crude extract and the purified lipopeptides, iturin and fengycin. We have selected 17 important plant and human pathogenic filamentous fungi (Table 1) ranging from the genera *Penicillium*, *Aspergillus*, *Talaromyces*, *Candida* to *Cryptococcus*. Besides those notorious plant pathogens such as *Aspergillus uvarum* infecting grapes, citrus rotting *Penicillium ulaiense* and post-harvest pathogen *Penicillium expansum* infecting apples, several important human pathogens were tested, such as *Aspergillus niger* and *Candida*.

The organic extracts of *B. velezensis* DTU001 displayed strong antifungal effects against *A. uvarum*, *P. ulaiense* and *P. expansum* (Table1 and Fig. 2). Due to the susceptibility to infection of mature and overripe fruit, post-harvest treatment of fruit with fungicides has traditionally been the most common method of combating *P. expansum*. Also, bio-fungicides using active ingredients such as bacteria and yeast have been successful in preventing infection but are ineffective against existing infections [33]. Considering the size of the apple product industry and the large number of people that may come into contact with infected fruits, control of the *P. expansum* is vitally important. Interestingly both *Talaromyces atroroseus* and *Talaromyces amestolkiae* were inhibited by both fengycin (20 mm) and iturin (29, 26 mm) as well as the crude extract (Table 1 and Fig. 2). All the tested organisms showed less or no susceptibility towards fengycin.

**Fig. 2.**
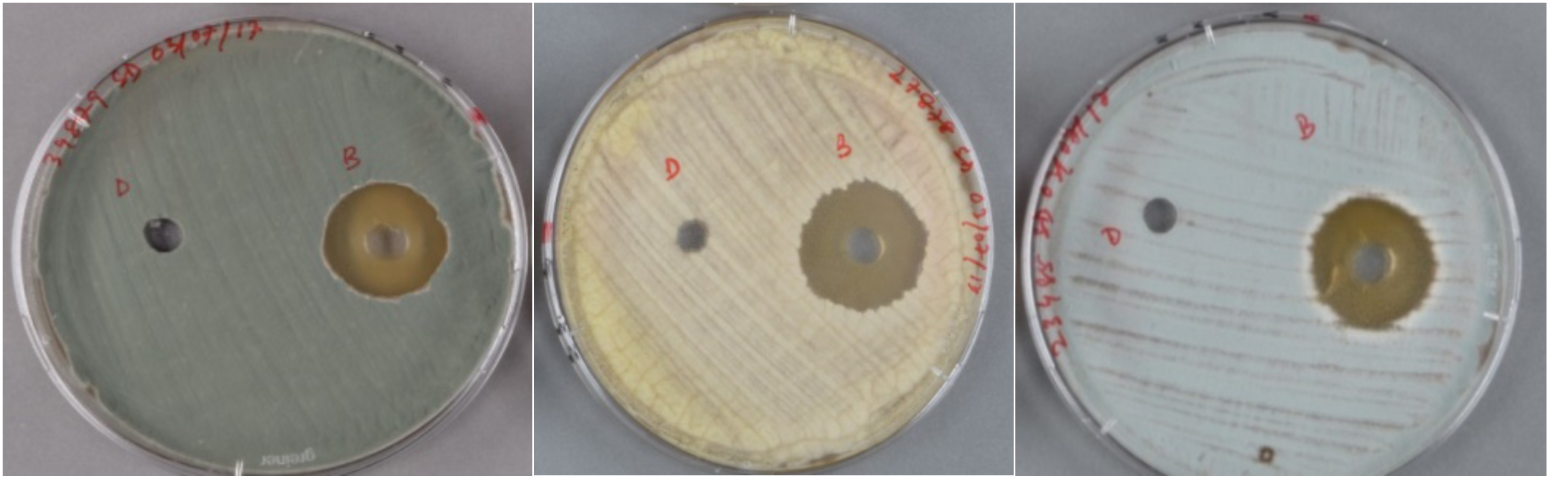
Extracts of *B. velezensis* DTU001 against fungi showing clear inhibition zones. Images left to right: *P. frequentans*, *A. niger* and *T. amestolkiae*. DMSO control loaded to the left hole, while the crude extract was applied to the right hole demonstrating the specific inhibition zone against the fungi.

In summary, the agar diffusion test showed that iturin and fengycin displayed selective anti-fungal activities against certain fungal pathogens, which means that some strains are insensitive to the addition of a single purified compound. For example, neither iturin nor fengycin displayed activities against *Penicillium sclerotigenum* and *Aspergillus calidoustus*. Overall, iturin displayed stronger and broader activities compared to fengycin. Compared to iturin and fengycin standards, extracts of *B. velezensis* DTU001 exhibited stronger inhibition activities and displayed a broader spectrum against test fungi including *P. sclerotigenum* and *A. calidoustus*. One explanation might be that *B. velezensis* produced a cocktail mixture which functions in a synergistic and more effective manner or additional secondary metabolite plays a role in the inhibition. However, the combined impact of the lipopeptides could avoid fast resistance evolution and help the bacterium in the environmental survival. It seemed that *Aspergillus felis* harbored certain mechanisms to avoid anti-fungal effects from *B. velezensis* DTU001 as neither lipopeptide standards nor the bacterial crude extract displayed inhibition activities against this fungus.

### 3.3. B. velezensis DTU001 is able to lessen biofilms of C. albicans

As *B. velezensis* DTU001 showed broad anti-fungal activity spectrum, we further evaluated if the bacterial isolate can directly reduce the growth of the human pathogen fungus, *C. albicans* under planktonic conditions and during biofilm development. First, the growth of the fungus and bacterium was monitored in planktonic co-cultures and compared to the mono-cultures (Fig. 3). Importantly, the *B. velezensis* cell count in the coculture with *C. albicans* did not differ significantly compared monoculture. In contrast, *C. albicans* planktonic cells between 6 and 24 hours diminished in the presence of *B. velezensis* DTU001, therefore the bacterial isolate was able to inhibit the proliferation of the fungus in the co-cultures. Similarly, *B. velezensis* DTU001 was able to lessen the biofilms of *C. albicans* when co-cultured. The inhibitory effect was even higher that under planktonic conditions after 24 hours (2-log decrease in fungal cell count). These experiments further support the applicability of *B. velezensis* DTU001 as biocontrol agent even in the presence of the target organism.

**Fig. 3.**
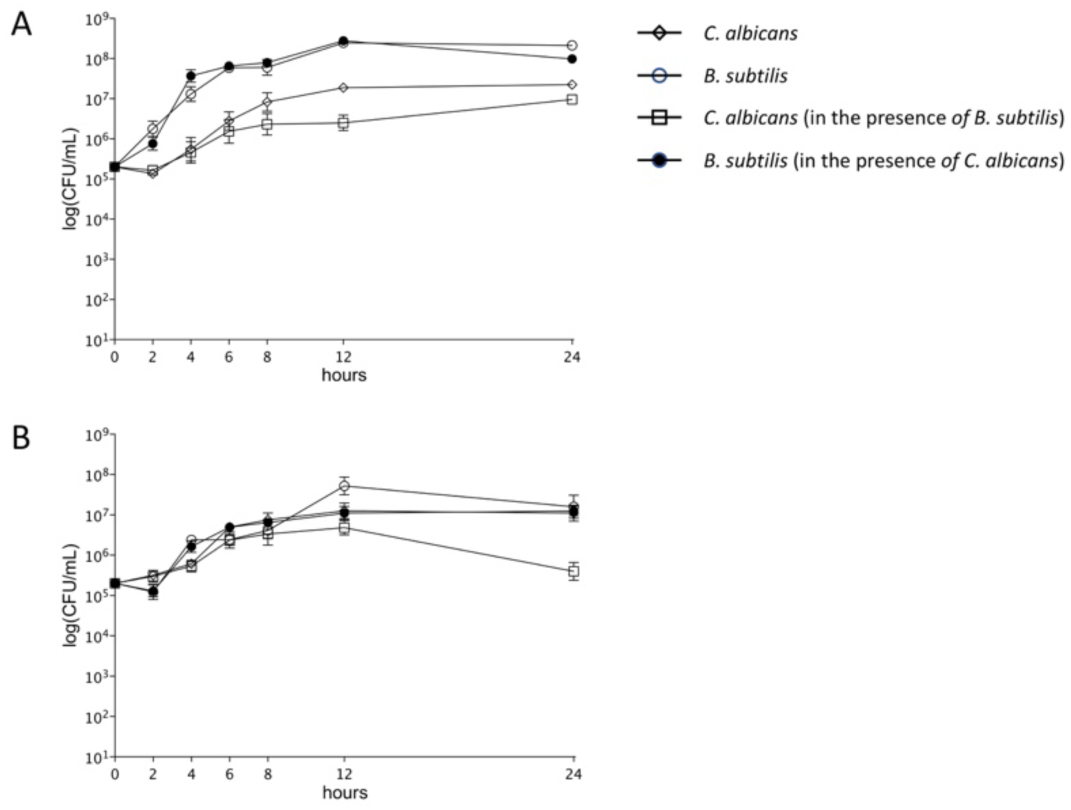
Planktonic (A) and biofilm (B) cultivation of *C. albicans* SC5314 and *B. velezensis* DTU001. Colony forming units were recorded at the indicated time points for *C. albicans* (open rhombus indicates *C. albicans* alone, open square denotes *C. albicans* in the presence of *B. subtilis*) and *B. velezensis* (open circle indicates *B. subtilis* alone, filled circle denotes *B. velezensis* in the presence of *C. albicans*).

### 3.4. B. velezensis DTU001 is a potent biofilm former

The probiotic effects of *Bacilli* depend on the ability to form biofilm and production of a matrix, that acts as an immune system stimulator [34], protects from environmental stress [35], and is required for plant colonization [36, 37]. Therefore, we have compared the biofilm formation of *B. velezensis* DTU001 to other *Bacillus* species known for biofilm formation. Both architecturally complex colonies on agar medium and air-liquid localized pellicles were tested. The biofilm development can be qualitatively monitored using these models: the more complex structures and enhanced wrinkles produced, the increased the biofilm development of *Bacilli* [28, 37]. In addition, pellicle formation strongly depends on the matrix components, without which cell are unable to float on the air-medium interface [8]. Both the architecturally complex biofilm colonies and the pellicles showed robust biofilm formation (i.e. wrinkled pellicles and colonies) by *B. velezensis* DTU001 that was comparable to other closely related isolates from the *B. subtilis* species complex (Fig. 4).

**Fig. 4.**
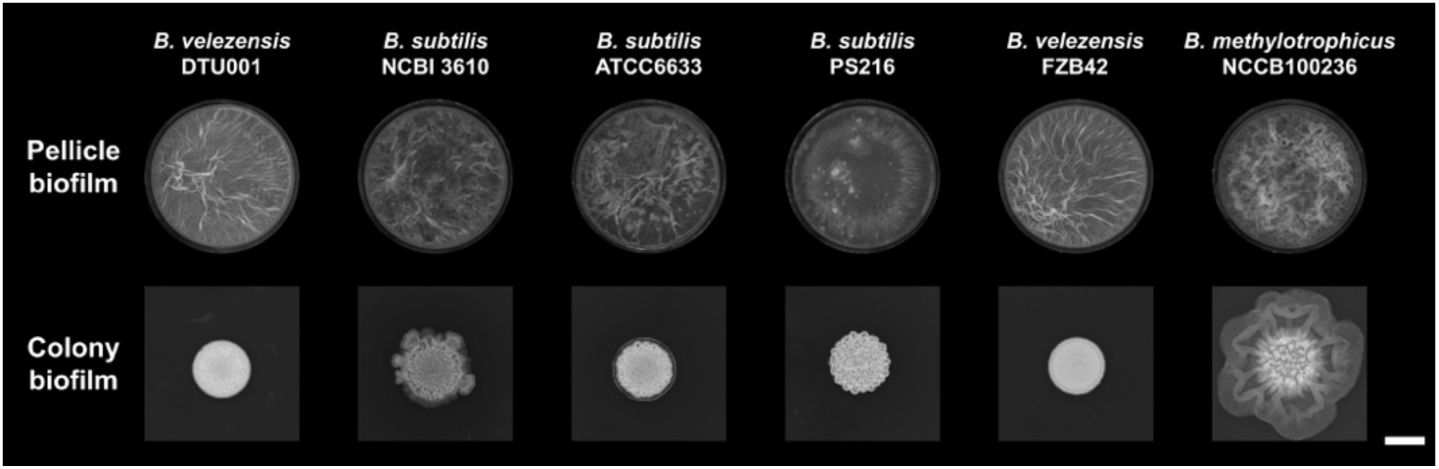
Pellicle (above) and colony biofilms (below) of selected strains from the *B. subtilis* species complex. The ability of biofilm formation is indicated by the development of wrinkles on the top of the pellicles and colonies. Scale bar indicates 5 mm.

### 3.5. Secondary metabolites characterization of B. velezensis DTU001

Based on the genome sequences and antifungal activity, *B. velezensis* DTU001 was predicted to produce various lipopeptides. To specifically demonstrate these secondary metabolites, LC-HRMS analysis was performed on ethyl acetate extracts harvested from agar grown *B. velezensis* DTU001. Indeed, chemical analysis along with purified standards detected three groups of bioactive lipopeptides: iturins, fengycins, and surfactins (Fig. 5, Fig. S1, S2, S3 and S4 in the supplemental data).

**Fig. 5.**
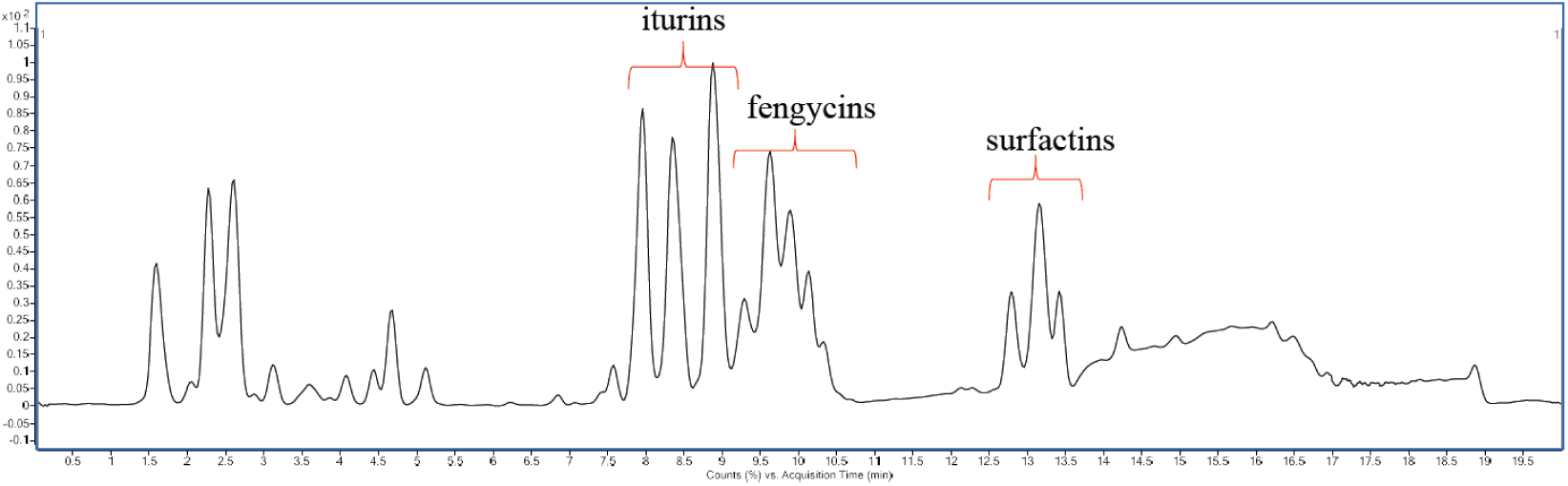
Base peak chromatogram showing the presence of iturin, fengycin and surfactin produced by *B. velezensis* DTU001.

## 4. Conclusions

Through genomics and metabolomics investigation, we demonstrated that *B. velezensis* DTU001 harbored the capacity to produce bioactive lipopeptides, comparable to the commercially utilized *B. amyloliquefaciens* and *B. velezensis* strains. Production of bioactive iturin, fengycin and surfactin by this strain was confirmed by HPLC-HRMS. Phylogenetic distance between DTU001 and those commercially utilized *B*. *amyloliquefaciens* together with their shared chemistry makes DTU001 as an attractive biocontrol option. Additionally, our targeted comparative genomics study revealed the versatility of secondary metabolite production by the selected *Bacilli*.

Either iturin or fengycin were previously proposed to be responsible for the antifungal activities of isolates within the *B. subtilis* species complex, however, we propose here that a cocktail anti-fungal agent derived from the crude extract of *B. velezensis* DTU001 functions more efficiently than single components. In addition, it will be interesting to examine whether purified surfactin also impacts the inhibitory potential of DTU001. In summary, this newly described isolate can be utilized as a biocontrol agent with a broad antifungal spectrum and provides a sustainable alternative for replacement of hazardous chemicals.

## Acknowledgements

This work was supported by the Danish National Research Foundation (DNRF137) for the Center for Microbial Secondary Metabolites (CeMiSt). SD was supported by a Novozymes and Henning Holck Larsen fellowship during her stay at DTU.

