## Supplementary data Fig S1 to S5 for "Depiction of secondary metabolites and antifungal activity of *Bacillus velezensis* DTU001"

### Supplementary Figures

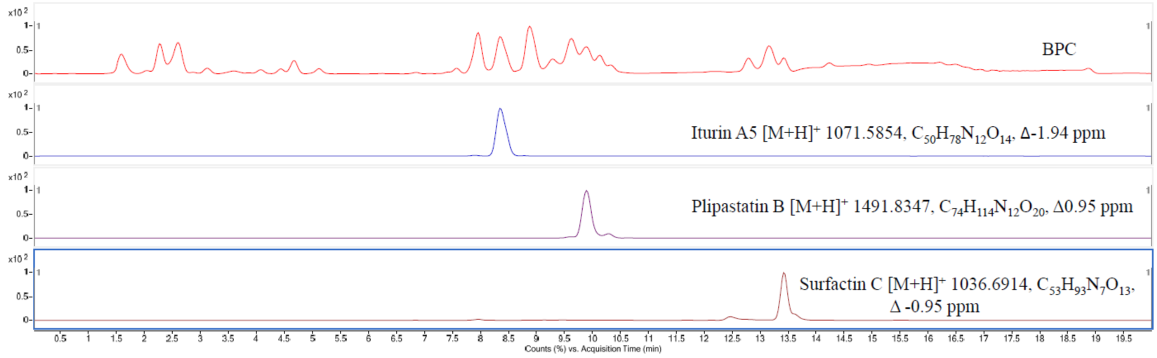


**Fig S1.** Detection of lipopeptides from *B. velezensis* DTU001 extract. Base Peak Chromatography of the crude extract (BPC, first line) and extracted ion chromatography indicating the presence of iturin A5 (second line), plipastatin B (third line) and surfactin C (fourth line).

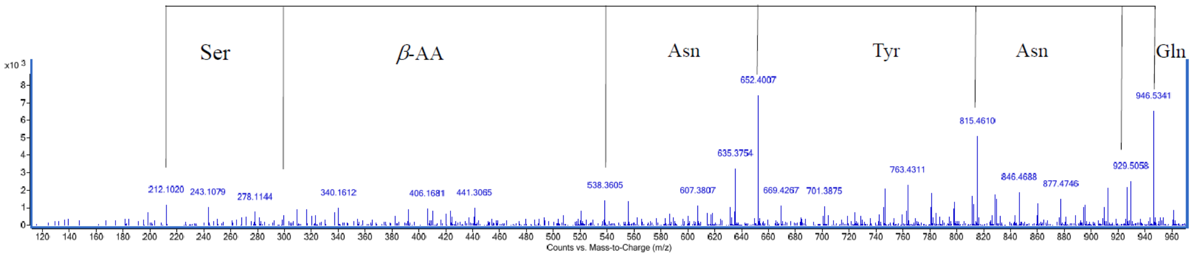


**Fig S2.** MS/MS fragmentation for iturin A5.

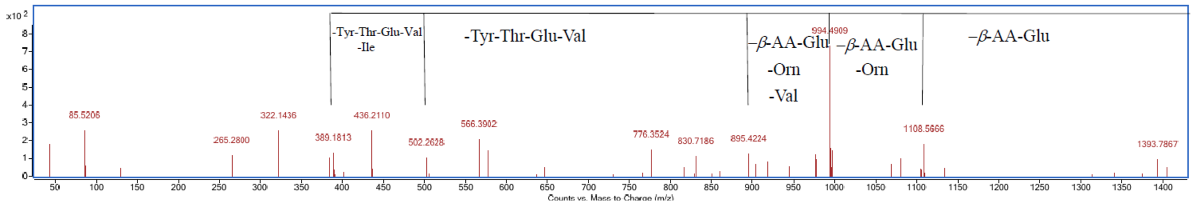


**Fig S3.** MS/MS fragmentation for plipastatin B.

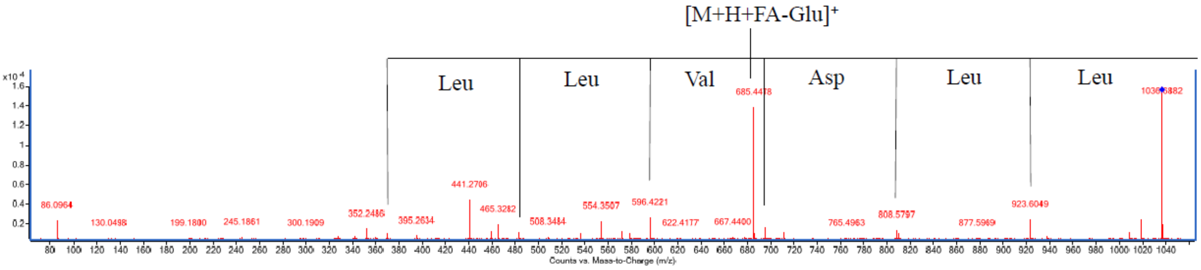


**Fig S4.** MS/MS fragmentation for surfactin C.
